# Maternal Gut Microbiota Regulates Neonatal Retinal Angiogenesis Through Lactational Metabolic Signalling

**DOI:** 10.64898/2026.05.29.728684

**Authors:** Sally A. Dreger, Raymond Kiu, Stephen D. Robinson

## Abstract

Early-life vascular development is a tightly regulated process that influences organ maturation and long-term cardiometabolic and neurovascular health. While the maternal microbiome is increasingly recognised as a determinant of neonatal immune, metabolic, and neurodevelopmental programming, its role in developmental angiogenesis remains poorly defined. Here, we investigated whether maternal gut microbiota regulates neonatal retinal vascular development through maternally derived metabolic signals.

Using antibiotic-induced dysbiosis with a broad-spectrum antibiotic cocktail consisting of vancomycin, neomycin, metronidazole, amphotericin B, and ampicillin (VNMAA), together with germ-free animals, we assessed retinal angiogenesis at postnatal day 6 (P6) and P12. Maternal microbiome depletion significantly delayed retinal vascular extension at P6 and produced persistent abnormalities in P12 retinal vascular architecture, including reduced deep plexus vascular density, altered descending branch density, and reduced branching complexity after normalisation to vessel density. Whole-genome shotgun sequencing demonstrated pronounced remodelling of the maternal caecal microbiome following antibiotic treatment and reciprocal faecal microbiota transplantation (FMT), whereas P12 pup caecal communities were comparatively limited and did not mirror the dramatic restructuring observed in dams. Restoration of the maternal microbiome by FMT rescued neonatal vascular defects, supporting a causal role for maternal microbial state.

Untargeted LC–MS metabolomics of maternal milk identified glycerophospholipid metabolism as a prominent microbiome-sensitive pathway. Choline and related lipid metabolites were prioritised as candidate mediators because of their position within this pathway and their biological relevance to membrane metabolism and endothelial function. *In vitro* endothelial assays demonstrated that choline modulates endothelial viability and promotes tubule formation.

Together, these findings identify a maternal microbiota–milk metabolite–vascular axis regulating neonatal retinal angiogenesis and suggest that lactational metabolic signalling may represent an important mechanism through which maternal microbial ecology shapes offspring vascular development.

## Introduction

Vascular development is a fundamental component of organogenesis and early postnatal tissue maturation. The formation, expansion, and remodelling of vascular networks ensure oxygen delivery, nutrient transport, immune surveillance, and metabolic homeostasis during periods of rapid growth. Disruption of vascular development during early life can therefore have consequences that extend beyond the immediate developmental window. This concept is consistent with the developmental origins of health and disease framework, which proposes that environmental exposures during critical windows of development can shape later physiological function and disease susceptibility^1–3^. While this framework has traditionally focused on maternal nutrition, placental insufficiency, hypoxia, and endocrine signalling, increasing evidence suggests that the maternal microbiome may represent an additional and underexplored determinant of offspring developmental programming.

Angiogenesis, the process by which new vessels sprout from pre-existing vasculature, is tightly regulated by growth factors, metabolic cues, extracellular matrix interactions, and endothelial cell-intrinsic signalling pathways^4^. The neonatal mouse retina provides a particularly powerful model for studying physiological angiogenesis because its vascularisation occurs postnatally in a stereotyped and temporally defined manner. During the first postnatal week, vessels expand radially from the optic nerve head to form the superficial vascular plexus. Subsequently, endothelial sprouts descend into the retina to form the deep and intermediate plexuses, generating a three-dimensional vascular network that supports the metabolic demands of the developing neural retina^5–8^. This predictable sequence allows investigators to distinguish early defects in vascular extension from later abnormalities in plexus formation, branching, and vascular maturation. The mouse retina is therefore widely used to study mechanisms of developmental angiogenesis and to model pathological retinal vascular disorders, including oxygen-induced retinopathy^7,9,10^.

Endothelial cells are highly responsive to their metabolic environment. During sprouting angiogenesis, endothelial cells must proliferate, migrate, extend filopodia, remodel membranes, and form lumenised networks. These processes require coordinated regulation of energy metabolism, lipid synthesis, membrane turnover, and redox balance. Canonical angiogenic pathways such as VEGF and Notch signalling are increasingly understood to act in concert with endothelial metabolic programmes^4,11,12^. For example, PFKFB3-driven glycolysis regulates endothelial tip-cell behaviour, cytoskeletal remodelling, and vessel branching, demonstrating that metabolic state is not merely supportive but functionally instructive during angiogenesis^11^. Although much attention has focused on glucose metabolism, lipid and phospholipid metabolism are also essential for endothelial proliferation, membrane biogenesis, and signalling. These observations raise the possibility that maternal or neonatal metabolic environments could directly influence angiogenic capacity during early development.

The gut microbiota is increasingly recognised as a regulator of host metabolism, immune maturation, and developmental physiology. In adults, microbial metabolites influence systemic metabolic tone, inflammatory pathways, and vascular function. In early life, the microbiome is particularly important because microbial colonisation occurs alongside rapid maturation of immune, metabolic, neurological, and epithelial systems. The infant gut microbiome is dynamic and undergoes functional maturation over the first months and years of life^13–15^. Maternal microbial communities contribute to this process through vertical transmission at birth, contact during nursing, and exposure to maternal-derived microbial products and metabolites. However, the neonatal gut microbiome is not immediately equivalent to the mature maternal microbiome; instead, it is sparse, dynamic, and shaped by delivery mode, feeding, antibiotics, and environmental exposure^13–16^. This distinction is important when interpreting early postnatal phenotypes: maternal microbial effects on offspring development may occur before the offspring microbiome is fully established.

A growing body of experimental work supports the idea that maternal microbiota can shape offspring development through mechanisms that do not require a mature offspring microbiome. Transient colonisation of germ-free pregnant mice demonstrated that the maternal microbiota can drive early postnatal innate immune development in offspring^17^. Maternal gut microbiota-derived short-chain fatty acids have been shown to influence offspring metabolic phenotypes through embryonic GPR41 and GPR43 signalling^18^. Maternal microbiome depletion and selective reconstitution can also alter foetal neurodevelopment in mice, with effects on axonogenesis and brain metabolite availability^19^. More recent metabolomic studies in germ-free and specific pathogen-free pregnancies have shown that maternal microbial status influences metabolite profiles in foetal intestine, brain, and placenta^20,21^. Together, these studies establish maternal microbiota as a developmental regulator acting through microbial metabolites, maternal physiology, and maternally transferred molecular cues.

Despite these advances, the role of maternal microbiota in vascular development remains poorly understood. Prior studies have implicated maternal microbial status in immune, metabolic, and neurodevelopmental outcomes, but whether maternal microbiota regulates developmental angiogenesis has not been defined. This represents a major gap because vascular development is essential for organ maturation and may represent an intermediary mechanism linking maternal exposures to long-term cardiometabolic and neurovascular risk. Moreover, because retinal vascularisation in mice occurs largely during the postnatal lactation period, it provides an opportunity to examine whether maternal microbial signals influence angiogenesis after birth through maternal-derived factors.

Breast milk is a biologically active interface between maternal physiology and neonatal development. Beyond macronutrient delivery, milk contains lipids, phospholipids, amino acids, oligosaccharides, immune mediators, hormones, microbial products, and small metabolites that contribute to immune maturation, microbial colonisation, growth, and organ development^22^. Human milk composition varies with gestational age, lactational stage, maternal metabolic status, and other maternal factors^22–26^. Metabolomic studies have shown that milk contains dynamic low-molecular-weight metabolite profiles and that these profiles are associated with infant growth and developmental outcomes^23–25^. Maternal adiposity and obesity have also been associated with changes in the human milk metabolome, supporting the concept that maternal physiology can alter milk-derived biochemical signals^27,28^. These observations make milk a plausible conduit through which maternal microbiome-dependent metabolic changes could influence neonatal vascular development.

Among milk-derived metabolic pathways, glycerophospholipid metabolism is particularly relevant to developmental biology and angiogenesis. Human milk phospholipids, including phosphatidylcholine and related lipid species, vary with gestational and lactational age and are recognised as bioactive components important for infant development^26^. Glycerophospholipids provide structural components for cellular membranes and act as precursors for signalling molecules involved in proliferation, migration, and cellular stress responses. Choline is a central metabolite within this pathway. It contributes to phosphatidylcholine synthesis, methyl-donor metabolism through betaine, acetylcholine production, and membrane biogenesis^29^. Choline availability is especially important during development, when rapid cellular proliferation and tissue growth increase demand for membrane synthesis and one-carbon metabolism^29,30^.

Choline also has precedent in developmental angiogenesis. Maternal dietary choline deficiency has been shown to alter angiogenesis in the foetal mouse hippocampus, decreasing endothelial cell proliferation and reducing vessel number while modifying expression of angiogenic genes such as VEGFc and Angpt2^31^. In endothelial cells, phosphatidylcholine metabolism is an active and regulated pathway, and exogenous phosphocholine can influence choline uptake and phosphatidylcholine synthesis^32^. These findings support the biological plausibility that altered choline or phosphatidylcholine-related metabolite availability could affect endothelial function during vascular development. However, whether maternal microbiota regulates milk choline-related metabolites, and whether these metabolites influence neonatal angiogenesis, has not been established.

Here, we test the hypothesis that maternal gut microbiota regulates neonatal retinal angiogenesis through microbiota-dependent changes in maternal milk metabolites. Using antibiotic-induced maternal dysbiosis, germ-free animals, reciprocal faecal microbiota transplantation, whole-genome shotgun sequencing, retinal vascular phenotyping, untargeted milk metabolomics, and endothelial functional assays, we define a maternal microbiota–milk metabolite–vascular axis. We show that maternal microbiome depletion delays early retinal vascular expansion and impairs deep plexus maturation, that restoration of the maternal microbiome rescues neonatal angiogenesis, and that maternal microbial state is associated with altered milk glycerophospholipid metabolism. Finally, we identify choline as a candidate microbiome-sensitive metabolite capable of modulating endothelial viability and angiogenic behaviour *in vitro*. Together, these findings expand the role of maternal microbiota in developmental programming to include regulation of neonatal vascular development.

## Materials and Methods

### Mice

Animal studies were designed and reported in line with ARRIVE 2.0 recommendations, with each individual pup considered the experimental unit. All animal procedures were conducted in accordance with the UK Animals (Scientific Procedures) Act 1986 and approved by the UK Home Office, in line with Directive 2010/63/EU on the protection of animals used for scientific purposes. C57BL/6J were purchased and maintained in-house at the Disease Modelling Unit (DMU), University of East Anglia, under project license code PP8873233 and under specific pathogen-free conditions with *ad libitum* access to food and water and appropriate environmental enrichment. Female mice were weighed upon setting up breeding trios and weighed weekly to determine gestation time. Once 2 weeks gestational, females were transferred to a new cage ready for treatment and weighed each time a treatment was administered.

Germ-free animals were maintained in sterile isolators under gnotobiotic conditions at the DMU. Germ-free pups were collected at matched developmental time points for retinal vascular analysis.

Animal experiments were designed in accordance with ARRIVE 2.0 recommendations. For *in vivo* experiments, the dam/litter was considered the unit of treatment allocation because microbiome depletion and FMT were applied at the maternal/litter level, while individual pups represented the unit of measurement for retinal vascular outcomes. Where multiple pups from the same litter were analysed, data are presented with biological repeat/litter structure indicated (colour coded), so that litter-to-litter variation can be visualised. Both male and female pups were included. Sex was recorded where possible but was not used as a stratification factor in the current study because experiments were not powered to detect sex-specific effects.

Sample sizes were based on previous experience with neonatal retinal angiogenesis assays and microbiome-manipulation studies, using the minimum number of animals required to detect biologically meaningful differences while accounting for litter-to-litter variability and maintaining ethical reduction of animal use. No formal *a priori* power calculation was performed. Exact n values for each analysis are reported in the relevant figure legends, where n indicates the number of individual retinas, pups, dams, wells, or biological repeats analysed, as appropriate. N indicates the number of independent litters, dams, or biological repeats, where applicable.

Where possible, pregnant dams were allocated to treatment groups to balance breeding date, litter age, and experimental processing order. For reciprocal FMT experiments, donor and recipient groups were processed in parallel to minimise confounding by collection date, cage location, staining batch, imaging batch, or analysis order. Germ-free animals were analysed as an independently maintained microbiota-deficient comparator group and were collected at matched developmental time points.

### Antibiotic-induced maternal microbiome depletion

Two-week gestational females were administered 0.2cc of a broad-spectrum antibiotic cocktail consisting of vancomycin, neomycin, metronidazole, amphotericin B, and ampicillin (VNMAA) by oral gavage three times weekly until the pups were born. Treatment continued to be administered to both the mothers and pups (pass the gavage needle over the pup’s lips) until the pups reached the age of P6 or P12. Control dams received regular drinking water.

The antibiotic cocktail contained vancomycin (5 mg/ml, Merck Life Science), metronidazole (10 mg/ml, Merck Life Science), neomycin (10 mg/ml, Merck Life Science), amphoteracin B (0.2 mg/ml, Merck Life Science) and ampicillin (1 mg/ml, Merck Life Science) prepared in drinking water.

Pups born to control and VNMAA-treated dams were collected at P6 or P12 for retinal vascular analysis. Maternal and pup caecal contents were collected for microbiome analysis in designated experiments.

### Faecal microbiota transplantation

Fecal pellets were collected separately (at P1) from both, control treated and antibiotic treated dams in sterile Eppendorf tubes and kept on ice. Pellets were weighed, a sterile solution of 10% glycerol (Merck) in normal saline (0.9% NaCl) was added to the pellets at a 30:70 fecal material to glycerol/normal saline solution, then centrifuged to produce a slurry which was stored at -80°C until use^33^.

Animal experiments were set up as above with treatment ceasing when pups were P6. At P7, recipient mothers were administered by oral gavage (0.2cc) with a reciprocal faecal microbiota transplantation (FMT). Experimental groups included control, VNMAA-treated, VNMAA dams receiving control FMT, and control dams receiving VNMAA-associated FMT. Retinas were collected from offspring at P12 for vascular phenotyping.

### Tissue collection

Pups were culled at P6 or P12. Eyes were enucleated and fixed in 4% paraformaldehyde for 1 hour at 4°C then transferred to 2X PBS for 30 minutes at 4°C. Retinas were dissected in phosphate-buffered saline (PBS) under a stereomicroscope, with care taken to remove the hyaloid vasculature and preserve retinal structure.

Maternal and pup caecal contents were collected aseptically at the time of sacrifice, snap-frozen in liquid nitrogen, and stored at −80°C until DNA extraction. Maternal milk was collected at P12 for metabolomic analysis as described below.

### Inclusion and exclusion criteria

The primary *in vivo* outcome measures were retinal vascular extension at P6 and P12, and P12 deep plexus vascular density. Secondary retinal outcome measures included superficial plexus vessel density, descending branch density, and branch-point measurements normalised to vessel density. For microbiome analyses, outcome measures included genus-level relative abundance profiles in maternal and pup caecal samples. For milk metabolomics, outcome measures included pathway enrichment and relative abundance of metabolites, with particular focus on glycerophospholipid metabolism and choline-related metabolites. For endothelial assays, outcome measures included HUVEC viability and HDF/HUVEC tubule formation.

Inclusion and exclusion criteria were defined before quantitative analysis. Pups were included if they were collected at the specified developmental stage, had intact eyes/retinas suitable for dissection, and belonged to the relevant experimental group. Retinas were included in plexus analysis only if staining quality, tissue integrity, and z-stack acquisition were sufficient to distinguish the superficial, intermediate/descending branch, and deep vascular layers. Images or regions were excluded only where technical artefacts prevented reliable quantification.

No animals or samples were excluded based on the observed vascular phenotype. Where exclusions occurred, they were due to technical failure during tissue collection, staining, imaging, or image analysis. Exact n values after exclusion are reported in the corresponding figure legends. Samples with insufficient sequencing depth or failed library preparation were excluded from microbiome analysis according to predefined quality-control thresholds.

### Retinal whole-mount immunostaining

Retinas (P6/P12) were stained for immunofluorescence as described previously by Benwell et al^34^. Briefly, after dissection, retinas were fixed for 20 minutes in ice cold methanol prior to blocking and immunostaining. Retinas were first permeabilized in 0.3% Triton/ PBS three times for ten minutes at room temperature (RT), washed twice (15 minutes each) in PBLEC in the dark, then blocked for 30 minutes in Dako Block (Agilent Dako #X090930-2) for 30 minutes in the dark at RT. Retinas were incubated overnight at 4°C in primary antibodies (made up in PBLEC) against isolectin B4/BS1-Lectin (clone L9831, Sigma 1:250). The retinas were washed three times for ten minutes in 0.1% Triton/ PBS in the dark at RT then mounted with Fluoromount (ThermoFisher). Retinas were imaged using a Zeiss LSM880 Airyscan Confocal microscope with identical acquisition settings across comparable groups. Whole-retina images were obtained for vascular extension analysis, while higher-magnification z-stacks images were obtained for superficial, intermediate (descending vessel), deep analyses. Vascular extension analysis (P6) and percentage of vascularization (P12) were performed using ImageJ while vessel density analysis of z-stacks (P12), was performed using Image first to convert images then analysed in AngioTool.

### Quantification of retinal vascular development

P6 vascular extension was quantified as the radial distance from the optic nerve head to the angiogenic front. Measurements were taken in three quadrants per retina and averaged to generate a single value per retina. P12 vascular extension was quantified similarly to assess whether the early delay in superficial vascular outgrowth persisted at later developmental stages.

For P12 plexus analysis, confocal z-stacks were separated into superficial, intermediate/descending branch, and deep vascular layers. Vessel density was quantified using ImageJ/Fiji and AngioTool following consistent image processing and thresholding across experimental groups. Vessel density was calculated for the superficial plexus, deep plexus, and descending branches and expressed as percentage relative to water-treated controls. Branching complexity was assessed by quantifying branch points and normalising branch-point measurements to vessel density, allowing branching changes to be interpreted independently of overall differences in vascular area.

To quantify average vascular density in the deep plexus at P12 (whole flat-mount), images were analysed using ImageJ/Fiji maintaining similar image processing and thresholding across the experimental groups. Vessel density was then expressed as a percentage relative to control.

Quantification was performed using identical image-processing and thresholding parameters across comparable experimental groups. Investigators performing image acquisition and quantitative image analysis were blinded to experimental group wherever possible. For retinal vascular extension and plexus analyses, the statistical measurement unit was an individual retina, while litter identity was retained as the biological repeat structure. Where data are shown as pooled N, each colour represents an independent biological repeat/litter and each point represents an individual retina within that biological repeat. For *in vitro* assays, each colour represents an independent biological repeat, with individual points representing technical wells or analysed fields as described in the figure legends.

### DNA extraction and whole-genome shotgun sequencing

Caecal material was weighed into MPBio Lysing Matrix E bead beating tubes (MPBiomedical, Germany) then DNA extraction was completed according to the manufacturer’s protocol for the MPBio FastDNA™ SPIN Kit for Soil (MPBiomedical, Germany) with an increase in the bead beating time to 3 minutes. DNA recovered from these samples was assessed using a Qubit® 2.0 fluorometer (Thermofisher).

Whole-genome shotgun sequencing was performed as described previously ^35^. Briefly, DNA was extracted from caecal material using the MPBio FastDNA SPIN Kit for Soil, with an extended bead-beating step. Sequencing libraries were prepared using a modified Illumina Nextera low-input tagmentation approach. Genomic DNA was normalised to 0.5 ng/µL, and 1 ng total DNA was used for tagmentation. Libraries were amplified using Nextera XT index primers, quantified, pooled in equimolar amounts, and size selected using KAPA Pure Beads. The final library pool was quantified using Qubit and assessed using an Agilent Tapestation prior to sequencing.

Libraries were sequenced on an Illumina NextSeq 500 using a Mid Output v2 300-cycle flow cell, with a 1% PhiX spike-in. Raw data were converted to FASTQ files using Illumina BaseSpace. FASTQ reads were quality-filtered (adapter sequenced removed) using fastp v0.21.0^36^. Host-associated reads were removed using Kraken2 v2.1.2^37^. Metagenomic purified reads were parsed into Kraken2 v2.1.2 for taxonomic profiling ((Kraken2 standard Refseq indexes retrieved from https://benlangmead.github.io/aws-indexes/k2), with confidence level set at 0.1. Bracken v2.6.2^38^ was then utilised to re-estimate the relative abundance of taxa at both genus and species level (-t set at 10) from Kraken2 outputs as recommended.

For functional analysis, clean reads were concatenated for each sample prior to analysis. Taxonomic profiling was performed using MetaPhlAn v3.0^39^ with the --unknown_estimation and --add_viruses options and the ChocoPhlAn v30 database. Functional profiling was performed using HUMAnN v3.0.0^40^ alpha with UniRef90 translated search mode and DIAMOND v0.9.24. Taxonomic and pathway outputs were normalised to relative abundance. Relative abundance data were analysed in R and visualised using ggplot2. Where applicable, differential taxa or pathway abundance was assessed using Wilcoxon–Mann–Whitney tests, with Benjamini–Hochberg correction for multiple testing.

### Maternal milk collection

Maternal milk was collected from lactating dams at P12. Dams were separated from pups for 4 hours prior to milk collection to allow milk accumulation. Dams were administered oxytocin (0.1cc (2 IU/kg)) intraperitoneally to stimulate milk let-down for 5 minutes then anaesthetised. Milk was collected manually from mammary glands using gentle pressure and capillary pipettes. After milk collection, dams were culled.

Milk samples were kept on ice for transport then stored at −80°C until metabolomic analysis. Samples were collected from control, VNMAA-treated, VNMAA + control FMT, and control + VNMAA FMT dams.

### Untargeted milk metabolomics

Untargeted metabolomic profiling of maternal milk (5-25μl of milk in a sterile Eppendorf tube) was performed by Biocrates Lifescience Ag (Innsburck, Austria) using the MxP® Quant 500 assay to measure endogenous metabolites. Flow injection analysis-tandem mass spectrometry (FIA-MS/MS) using a SCIEX API 5500 QTRAP was used to measure lipids and hexose, while liquid chromatography-tandem mass spectrometry (LC-MS/MS) was used to measure small molecules. Sciex Analyst® software was used to calculate metabolite concentrations and biocrates’ MetIDQ™software was used for further data analysis. MetaboAnalyst (free online software) was used to analyze the returned Biocrates metabolomic data.

Pathway enrichment analysis was performed to identify metabolic pathways altered across experimental groups. Comparisons included control versus VNMAA and reciprocal FMT conditions. Particular attention was given to glycerophospholipid metabolism and associated metabolites, including choline, serine, methionine, total diacylglycerides, total lysophosphatidylcholine, and total phosphatidylcholine.

### Endothelial cell culture

Human umbilical vein endothelial cells (HUVECs, C-12203, Merck) were cultured in M199 media (Merck) supplemented with 20% foetal bovine serum (FBS), 1% penicillin-streptomycin, 1% L-glutamine, 0.1mg/ml Endothelial Cell Growth Supplement, 0.1mg/ml heparin (Merck) in a humidified incubator at 37°C with 5% CO₂. Cells were used between passages (P7-P11) and were routinely monitored for morphology and confluence.

For co-culture tubule formation assays, HUVECs were cultured with human dermal fibroblasts (HDFs, C-12302, Merck). HDFs were cultured in high glucose DMEM with 10% FBS, 1% penicillin-streptomycin, 1% L-glutamine (Merck) at the same conditions as above. Cells were treated with choline at concentrations ranging from 0µM to 20µM. VEGF at 30ng/mL served as a positive pro-angiogenic control, and suramin at 30µM served as a negative control.

To assess both endothelial cell viability and co-culture tubule formation assays, cells were grown in serum free/Choline Free IMMLEC media (CFM) prepared using Dulbecco’s MEM (DMEM) High Glucose, w/ L-Glutamine, Pyridoxine HCl, Sodium Bicarbonate, w/o Glycine, Serine, Choline, Chloride (D9800-14-25L, Generon) supplemented with varying concentrations of choline chloride (C7017-5G, Merck) made from a 100mM stock solution.

### Endothelial viability assay

HUVEC viability was assessed following treatment with varying concentrations of choline. In brief, 96 well-plates were coated with 10µg/ml of fibronectin (FN, F0895-5MG, Merck), incubated for 30 minutes then FN was removed. HUVECs were seeded at a density of 2x10^4^ cells/well, incubated overnight at 37°C, 5% CO_2_ in 150μl/well of complete IMMLEC media (contains 10% FBS). After 24 hours, media was removed and wells washed twice with sterile PBS. Cells were serum starved for 3 hours in 150μl/well of serum free/Choline Free IMMLEC Media (CFM), media was removed then 100µl of assay medium (Choline Free IMMLEC (CFM) +/- Choline concentrations (0µM, 0.1µM, 0.5µM, 1µM, 2.5µM, 5µM, 10µM, 15µM, 20µM); complete IMMLEC; 30ng/ml VEGF in CFM; 30μM Suramin in CFM) in columns of eight and incubated for 24 hours. Endothelial cell viability/number was assessed using the CellTiter 96^®^ Aq_ueous_ One Solution Cell Proliferation Assay (Promega, Madison, WI, USA) according to manufacturer’s instructions. Warmed reagent was prepared 1:5 in choline free media, then 120µl was added directly to culture wells containing HUVEC cells, after incubation for 1 hour at 37°C, plates were read at A_492_ using a 96-well plate reader read on a microplate reader (VersaMax, Molceular Devices, USA) whereby absorbance is directly proportional to the number of viable cells. Optical density values were normalized to untreated control or vehicle-treated cells and expressed as percentage viability relative to control with graphs generated in Prism 10.

### Tubule formation assay

Endothelial tubule formation was assessed using an HDF-HUVEC co-culture assay. Briefly, 24-well plates were coated with FN as above, then HDFs were seeded at a density of 5 x 10^4^ cells/well and incubated in HDF media at 37°C/ 5% CO_2_ for 72 hours. Next HUVECs (1 x 10^4^ cells/well) were seeded onto the HDF layer in either, assay medium (Choline Free IMMLEC (CFM) +/- Choline concentrations (see above); complete IMMLEC; 30ng/ml VEGF in CFM; 30μM Suramin in CFM) in triplicate. Plates were Incubated at 37°C/ 5% CO_2_ (Day 1) then fed as above on Days 3 and 5. On Day 8, wells were fixed with ice cold 70% ethanol for 30 minutes at room temperature, washed three times for 5 minutes with PBS then incubated with mouse anti-human CD31 primary antibody (BioRad, MCA1738) for 1 hour at 37°C. Wells were washed again with PBS (as above) then incubated with goat anti-mouse IgG1 secondary antibody (BioRad, STAR132A) for 1 hour at 37°C followed with another wash with distilled water (dH_2_O), 3 times for 5 minutes. Finally, 1 BCIP/NBT tablet (#B5655, Merck) was added to 10ml of water, filtered with a 0.2μm filter, added to wells, incubated at 37°C for 15 minutes. (do not exceed 30min) then washed with dH_2_O and air-dried. Images were taken on a SP-105 Inverted Fluorescent microscope (Brunel Microscopes Ltd), then tubule formation was quantified using ImageJ/AngioTool to assess the percentage tubule formation, total tubule length and number of branch points. Quantification was performed on 3 fields per well and 3 technical replicates per condition with graphs and statistics generated in Prism 10 (GraphPad).

### Statistical analysis

Data are presented as mean ± SEM unless otherwise stated. Statistical analyses were performed using GraphPad Prism version 10 and R where indicated for microbiome or metabolomics analyses. For comparisons between more than two experimental groups, one-way ANOVA was used when data were approximately normally distributed and variance was comparable between groups, followed by appropriate post hoc multiple-comparison testing. Where these assumptions were not met, non-parametric alternatives were used. For experiments involving more than one independent variable, two-way ANOVA with appropriate post hoc correction was used where applicable.

For retinal vascular analyses, each individual retina was treated as the measurement unit, with litter or biological repeat structure shown. Plots show pooled biological repeat/litter effects, with each colour representing an independent biological repeat and each point representing an individual retina, well, or field within that repeat, as specified in the figure legends. For *in vitro* endothelial assays, data were generated from independent biological repeats with technical replicate wells or fields nested within each repeat.

For microbiome analyses, taxonomic relative abundance data were analysed in R. Where beta-diversity analysis was performed, differences in community structure were assessed using PERMANOVA. For metabolomics pathway analysis, p-values were corrected for multiple testing where applicable using false-discovery-rate correction or the provider-recommended method. Exact statistical tests and significance thresholds are indicated in the figure legends or on the graphs. P < 0.05 was considered statistically significant.

## Results

### Maternal microbiome depletion impairs early postnatal retinal angiogenesis

To determine whether maternal microbiota influences neonatal vascular development, we first depleted the maternal microbiome using VNMAA administered across the perinatal period, with retinal angiogenesis assessed at P6 and P12. Germ-free animals were included as an independent microbiota-deficient comparator (Figure 1A, schematic).

**Figure 1.**
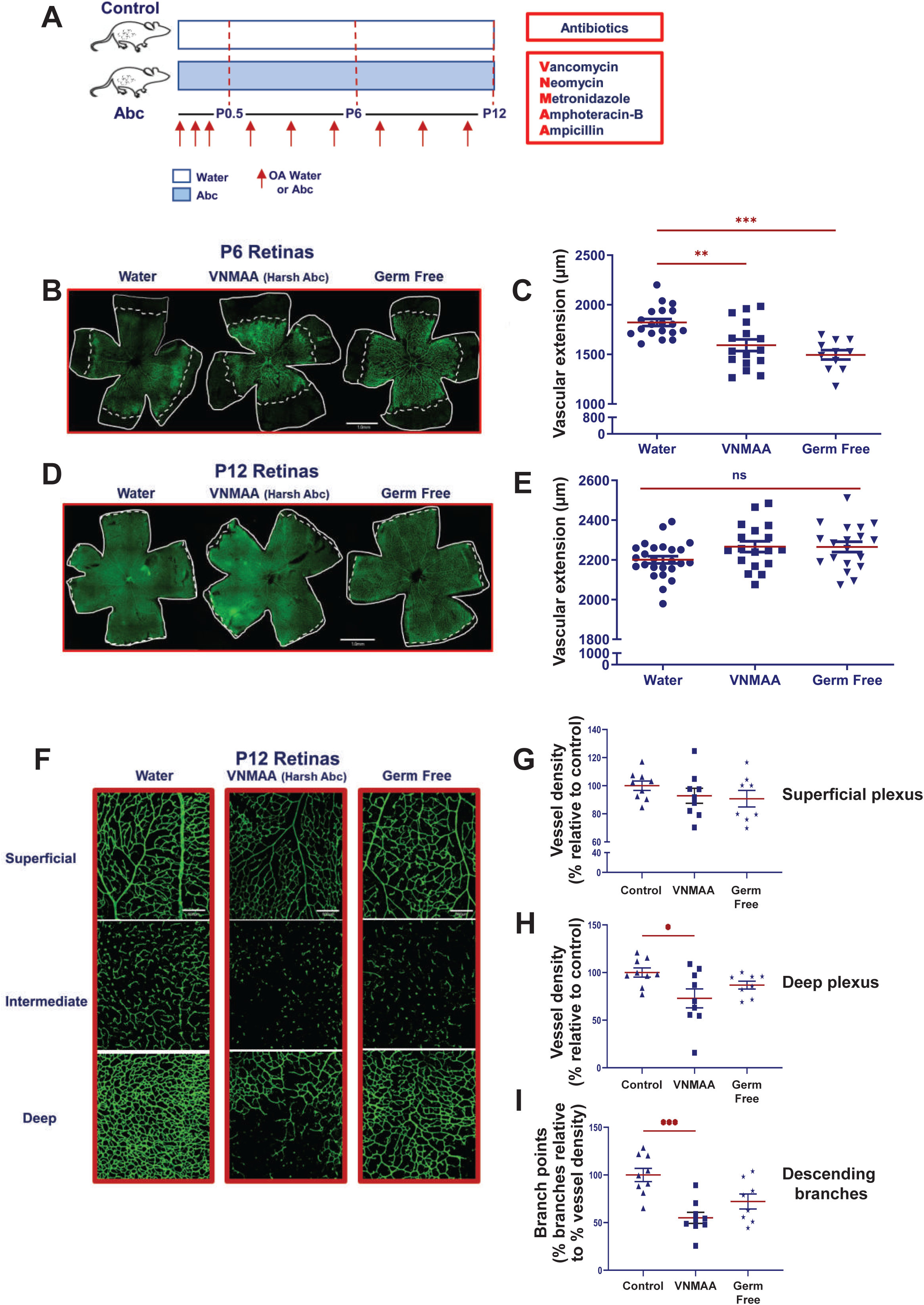
Early-life microbiome depletion transiently delays retinal vascular extension and alters retinal vascular plexus development. (A) Experimental design. Two-week gestational dams were treated with either water or a broad-spectrum antibiotic cocktail by oral gavage to induce gut dysbiosis. Neonatal mice were treated from P0.5 to P12 with water or antibiotics. Antibiotic-treated mice received VNMAA, consisting of vancomycin, neomycin, metronidazole, amphotericin-B, and ampicillin. Retinas were collected at P6 and P12. Germ-free mice were analysed in parallel as an independent microbiome-depleted condition. Red arrows indicate oral administration of water or antibiotics. (B) Representative BS-1-labelled whole-mount P6 retinas from water-treated control, VNMAA-treated, and germ-free mice. Retinal vasculature is shown in green. White outlines indicate the retinal edge, and dashed white lines indicate the vascular front used to measure radial vascular extension. Scale bar, 1.0mm. (C) Quantification of P6 vascular extension, measured as radial distance from the optic nerve head to the vascular front. VNMAA-treated and germ-free mice showed reduced vascular extension compared with water-treated controls. Each point represents an individual retina. Red horizontal lines indicate mean ± SEM. n = 11–19 retinas from N = 3 independent litters per group. (D) Representative BS-1-labelled whole-mount P12 retinas from water-treated control, VNMAA-treated, and germ-free mice. Retinal vasculature is shown in green. White outlines indicate the retinal edge. Scale bar, 1.0mm. (E) Quantification of P12 vascular extension. By P12, radial vascular extension was not significantly different among water-treated, VNMAA-treated, and germ-free mice, indicating recovery of superficial vascular outgrowth. Each point represents an individual retina. Red horizontal lines indicate mean ± SEM. n = 18–25 retinas from N = 3 independent litters per group. ns, not significant. (F) Representative confocal z-stack images of BS-1-labelled P12 retinal vasculature in water-treated control, VNMAA-treated, and germ-free mice. Images show the superficial, intermediate/descending branch, and deep vascular layers. Green signal indicates BS-1-positive retinal vessels. Scale bar, 500µm. (G) Quantification of P12 retinal vascular architecture in control, VNMAA-treated, and germ-free mice. Vessel density was quantified in the superficial plexus, deep plexus, and descending branches and expressed as percentage relative to water-treated controls. Branch-point measurements were normalised to vessel density and expressed as percentage fold change relative to control, allowing branching complexity to be assessed independently of total vessel coverage. Each point represents an individual analysed retina or field, as indicated by the source data structure. Red horizontal lines indicate mean ± SEM. For plexus analysis, n = 3 mice per group from N = 3 independent litters per group. Statistical significance is indicated on the graphs: *P < 0.05, **P < 0.01, ***P < 0.001. For all retinal analyses, each point represents an individual retina. Where multiple pups were analysed from the same litter, litter identity was retained as the biological repeat structure. Quantification was performed blinded to experimental group wherever possible.

At P6, pups born to VNMAA-treated dams exhibited reduced radial vascular outgrowth compared with controls, consistent with delayed expansion of the superficial retinal vascular plexus. Representative retinal whole-mount images showed a shorter distance between the optic nerve head and angiogenic front in microbiome-depleted groups (Figure 1B). Quantification confirmed a significant reduction in vascular extension in both VNMAA-exposed and germ-free animals compared with water-treated controls (Figure 1C). By contrast, total vascular extension at P12 was no longer significantly different between groups, suggesting that the earliest effect of microbiome depletion is a delay in developmental progression rather than a complete arrest of vascular growth (Figure 1D,E).

We next asked whether this apparent recovery in radial extension was accompanied by persistent abnormalities in retinal vascular architecture. Confocal imaging of P12 retinas across the superficial, intermediate/descending branch, and deep vascular layers revealed that microbiome-depleted animals showed the most pronounced abnormalities in the deeper vascular compartments (Figure 1F). Quantitative analysis showed that superficial plexus vessel density was relatively modestly affected, whereas deep plexus vessel density was reduced in microbiome-depleted animals (Figure 1G). Descending branch density was also reduced, consistent with impaired vertical sprouting and delayed formation of the deeper retinal vascular network (Figure 1G). In addition, branch-point analysis normalised to vessel density indicated altered vascular complexity, particularly within the deep plexus, suggesting that microbiome depletion affects not only the amount of vascular coverage but also the organisation of the developing vascular network (Figure 1G).

Together, these findings indicate that loss of maternal microbiota transiently delays early superficial retinal vascular expansion and leads to persistent defects in P12 retinal vascular architecture, particularly affecting deep plexus maturation, descending branch formation, and branching complexity.

### Restoration of the maternal microbiome rescues neonatal angiogenesis

The retinal phenotype raised a key mechanistic question: was impaired angiogenesis caused by microbiota depletion itself, and could this phenotype be reversed by restoring the maternal microbial community? To test this, we performed reciprocal FMT experiments during the postnatal period. VNMAA-treated dams received microbiota from control donors, while control dams received microbiota from VNMAA-treated donors (Figure 2A, schematic). Retinal vascular development was then assessed in offspring at P12, with particular focus on the deep vascular plexus, which was the compartment most sensitive to microbiome depletion (see Figure 1).

**Figure 2.**
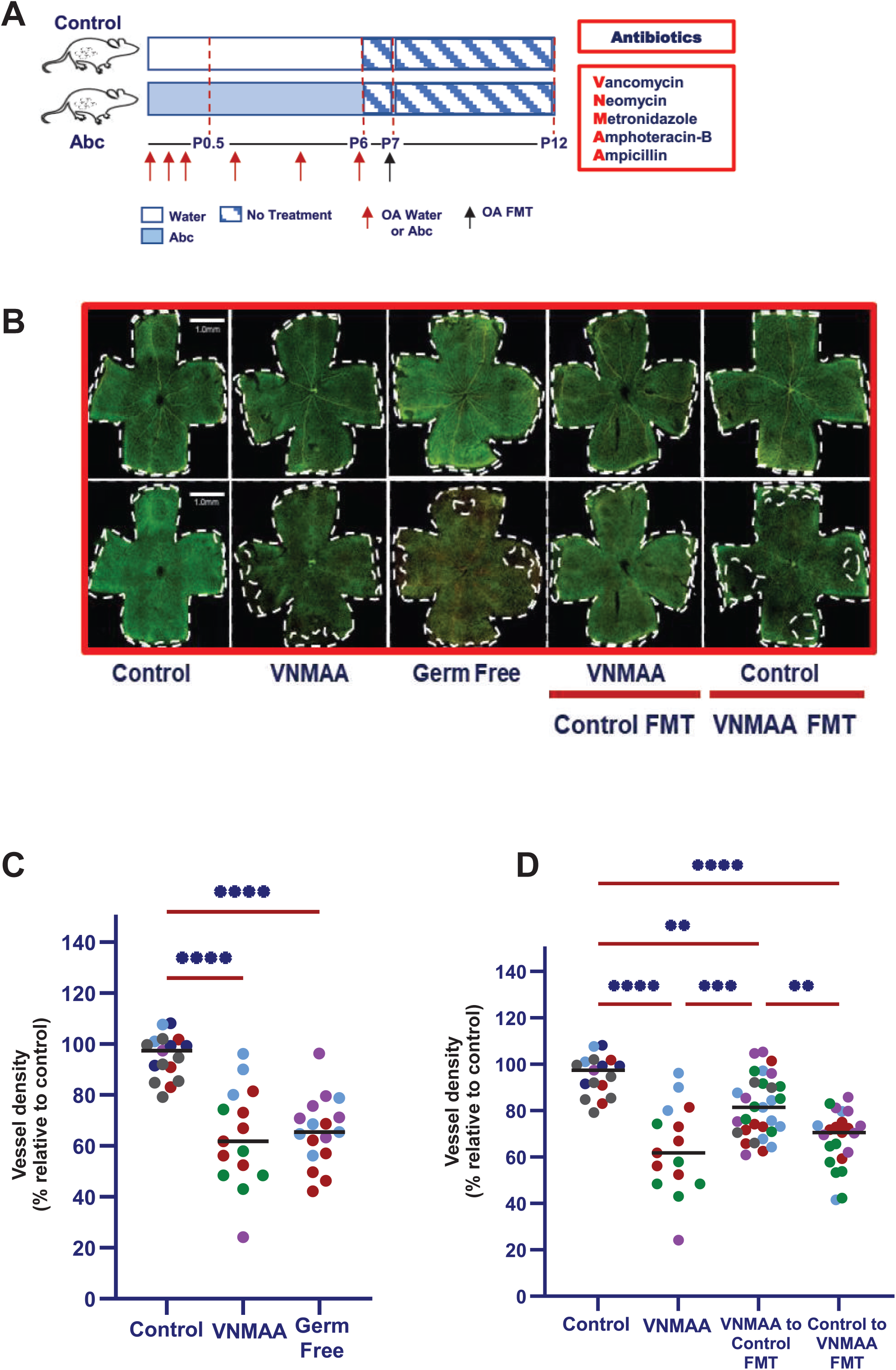
Restoration of the maternal microbiota rescues neonatal retinal angiogenesis. (A) Experimental design. Two-week gestational dams were treated with either water (control) or broad-spectrum antibiotic cocktail by oral gavage while, neonatal mice were treated from P0.5 to P6 with water or broad-spectrum antibiotics. Antibiotic-treated mice (Abc) received VNMAA, consisting of vancomycin, neomycin, metronidazole, amphotericin-B, and ampicillin. From P6 to P12, mice received no treatment. Reciprocal FMT was performed at P7, with VNMAA-treated dams receiving control FMT and control dams receiving VNMAA FMT. Retinas were collected at P12. Germ-free mice were analysed in parallel as an independent microbiome-depleted condition. Red arrows indicate oral administration of water or antibiotics, and the black arrow indicates oral FMT. (B) Representative BS-1-labeled P12 retinal whole-mount images from control, VNMAA-treated, germ-free, VNMAA-treated plus control FMT, and control plus VNMAA FMT groups. Retinal vasculature is shown in green. Images show superficial and deep vascular plexus preparations. White outlines indicate the retinal edge, and dashed white lines indicate the vascular front/vascularised region. Scale bar, 1.0mm. (C) Quantification of average vascular density in the deep plexus at P12, expressed as percentage relative to control. VNMAA-treated and germ-free mice showed reduced vascular density compared with controls. Data are presented as superplots, where each colour represents an independent biological repeat/litter and each point represents an individual retina within that biological repeat. Black horizontal lines indicate mean ± SEM. n = 15-17 retinas from N = 3-4 independent litters per group. (D) Quantification of average vascular density in the deep plexus at P12 following reciprocal FMT, expressed as percentage relative to control. VNMAA treatment reduced vascular density compared with controls, whereas transfer of control microbiota into VNMAA-treated dams partially restored vascular density. Conversely, transfer of VNMAA-associated microbiota into control dams reduced vascular density relative to controls. Data are presented as superplots, where each colour represents an independent biological repeat/litter and each point represents an individual retina within that biological repeat. Black horizontal lines indicate mean ± SEM. n = 15–31 retinas from N = 4-5 independent litters per group. Data are presented as pooled N: each colour represents an independent biological repeat, and each point represents an individual retina within that biological repeat. The black horizontal line indicates the group mean. Quantification was performed blinded to experimental group wherever possible. Statistical significance is indicated on the graphs: **P < 0.01, ***P < 0.001, ****P < 0.0001.

Representative P12 retinal whole-mount images showed that pups from VNMAA-treated dams had reduced vascular density, most prominently in the deep plexus. Germ-free animals showed a similar impairment, reinforcing the conclusion that the phenotype is linked to microbiota depletion rather than antibiotic exposure alone. In contrast, pups from VNMAA-treated dams that received control FMT showed visibly improved vascularisation, with deep plexus architecture more closely resembling control animals (Figure 2B).

Quantification confirmed this pattern. As previously, average vascular density was significantly reduced in VNMAA and germ-free groups compared with controls (Figure 2C). However, restoration of control microbiota to VNMAA-treated dams increased vascular density toward control levels. Conversely, transfer of VNMAA-associated microbiota into control dams shifted the offspring phenotype toward reduced vascular density (Figure 2D).

These reciprocal effects demonstrate that maternal microbial state is sufficient to modulate neonatal retinal angiogenesis. However, they did not yet distinguish whether the rescue reflected altered microbial colonisation of the pups themselves or indirect effects of the maternal microbiome on the neonatal environment. This distinction was important because pups remain highly dependent on maternal inputs during this developmental window.

### Maternal microbiome remodelling, rather than pup caecal colonisation, redirects the mechanistic focus toward maternal milk

To distinguish whether the retinal phenotype was associated with altered microbial colonisation of pups or with changes in the maternal microbiome, we performed whole-genome shotgun sequencing of maternal and P12 pup caecal samples across control, VNMAA, and reciprocal FMT groups.

The maternal microbiome showed a striking response to experimental manipulation. Control dams harboured a diverse microbial community, whereas VNMAA-treated dams exhibited a markedly altered genus-level profile. FMT further reshaped maternal microbial composition, demonstrating that microbial community structure could be redirected by transfer of control or dysbiotic material. Thus, the maternal compartment showed clear treatment-dependent microbial remodelling (Figure 3A).

**Figure 3.**
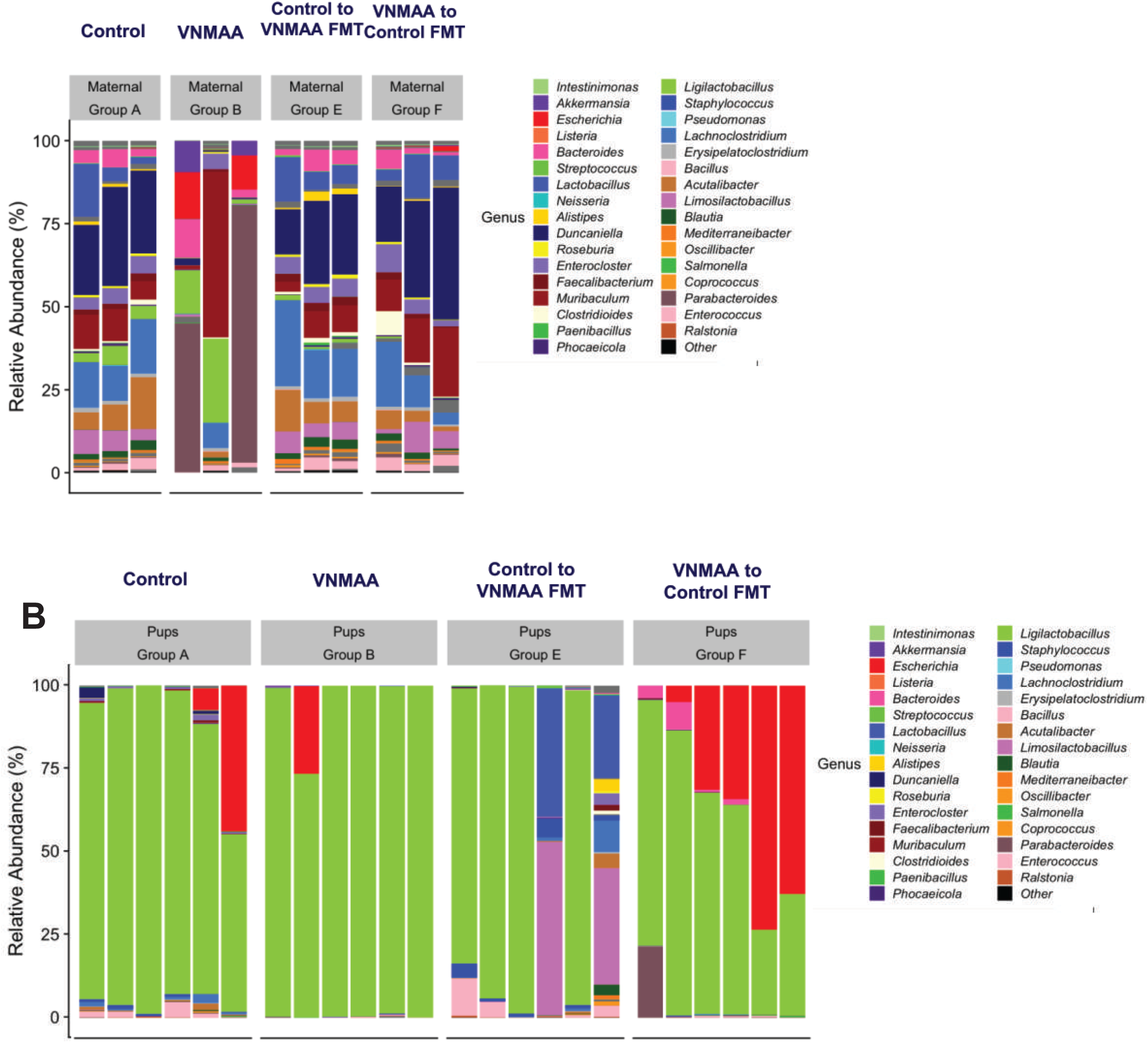
Maternal microbiome remodelling, rather than pup caecal colonisation, redirects the mechanistic focus toward maternal-derived signals. (A) Genus-level relative abundance profiles from whole-genome shotgun sequencing of maternal caecal samples. Samples are grouped as control, VNMAA-treated, control to VNMAA FMT, and VNMAA to control FMT. Each stacked bar represents an individual maternal caecal sample. Colours indicate bacterial genera, with low-abundance or unclassified taxa grouped as “Other.” Control dams showed a diverse genus-level microbial community, whereas VNMAA-treated dams showed marked restructuring of the maternal microbiome. Reciprocal FMT further altered maternal microbial composition, indicating treatment-dependent remodelling of the maternal microbial compartment. (B) Genus-level relative abundance profiles from whole-genome shotgun sequencing of P12 pup caecal samples from the same experimental groups. Each stacked bar represents an individual pup caecal sample. Colours indicate bacterial genera, with low-abundance or unclassified taxa grouped as “Other.” In contrast to the maternal caecal microbiome, P12 pup caecal communities were comparatively limited and were dominated by a small number of genera, particularly *Intestinimonas* and, in some samples, *Escherichia*. Pup profiles did not recapitulate the broad treatment-dependent restructuring observed in dams, suggesting that the retinal vascular phenotype is unlikely to be explained primarily by establishment of a mature, treatment-patterned pup gut microbiome.

By contrast, the P12 pup caecal microbiome did not show the same pattern. Pup samples contained comparatively limited microbial profiles and did not recapitulate the dramatic community shifts seen in the dams (Figure 3B). Although some differences were detectable across pup groups, these profiles did not provide a compelling explanation for the magnitude or direction of the retinal vascular phenotype.

This distinction was central to our interpretation. The vascular phenotype occurred during a developmental window when the pup gut microbiome remained relatively immature, while the maternal microbiome was robustly altered by both antibiotic treatment and FMT. We therefore reasoned that the maternal microbiome was likely acting through maternal-derived signals rather than through direct establishment of a mature offspring microbiome.

Because neonatal pups are dependent on maternal milk throughout this period, we next investigated whether microbiome-dependent changes in the milk metabolome could mediate the observed effects on retinal angiogenesis.

### Maternal microbial state reshapes the breast milk metabolome

The FMT experiments demonstrated that altering the maternal microbiome was sufficient to modify neonatal retinal angiogenesis. However, because P12 pup caecal communities were comparatively limited and did not mirror the dramatic microbial remodelling observed in dams, we reasoned that the maternal microbiome may influence offspring vascular development through maternally derived factors rather than through direct establishment of a mature pup gut microbiome. Given the dependence of neonatal pups on maternal milk during this developmental window, we next examined whether maternal microbial state was associated with changes in milk metabolite composition.

To test this, we performed untargeted LC–MS metabolomic profiling of maternal milk collected at P12 from control, VNMAA-treated, and reciprocal FMT dams. Pathway enrichment analysis revealed broad metabolic differences between control and VNMAA milk samples, with glycerophospholipid metabolism emerging as one of the most prominently enriched pathways. Additional altered pathways included glutathione metabolism, D-glutamine and D-glutamate metabolism, pantothenate and CoA biosynthesis, nitrogen metabolism, arginine and proline metabolism, tryptophan metabolism, arachidonic acid metabolism, glycerolipid metabolism, primary bile acid biosynthesis, and biosynthesis of unsaturated fatty acids (Figure 4A).

**Figure 4.**
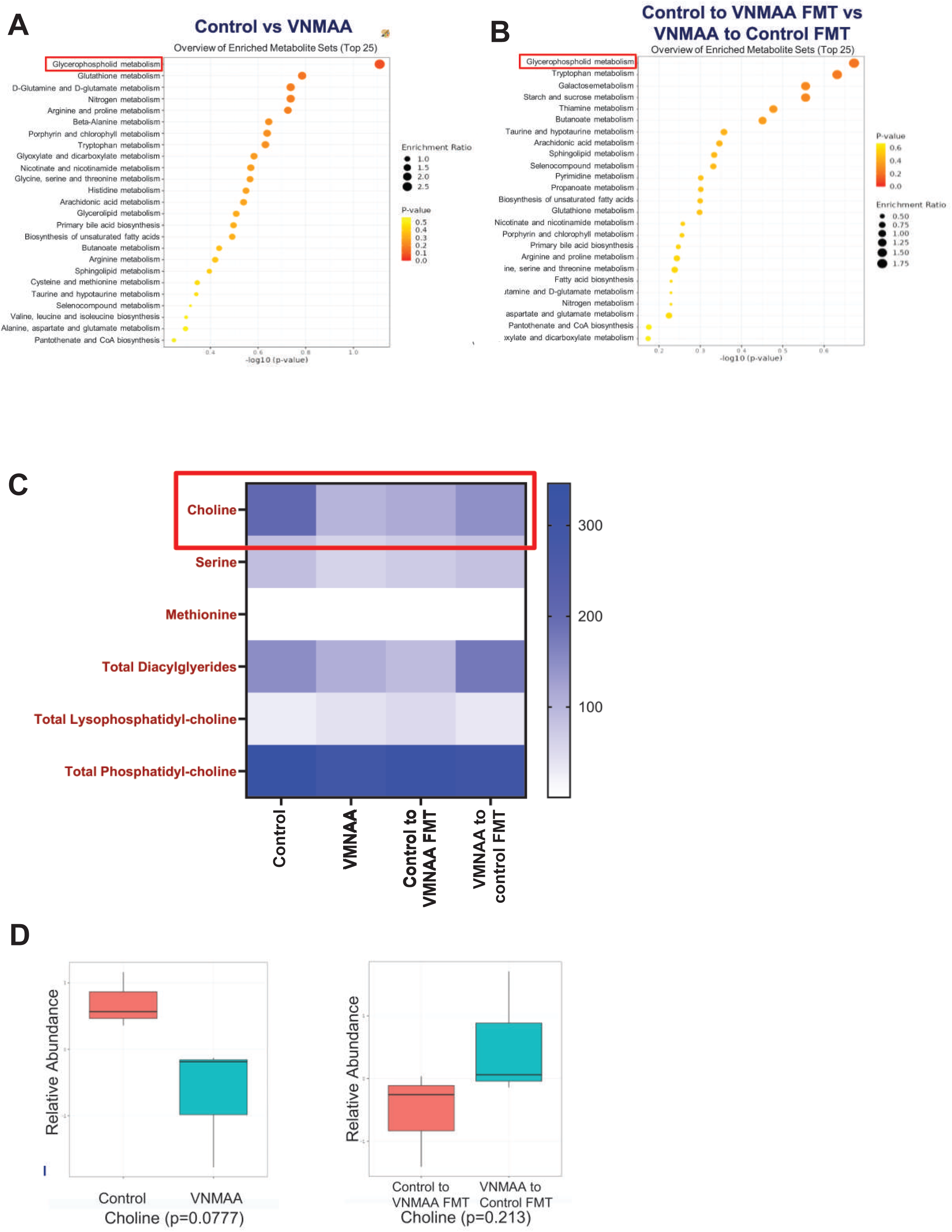
Maternal milk metabolomics identifies glycerophospholipid metabolism and choline as candidate microbiome-dependent pathways linked to neonatal vascularisation. (A) Pathway enrichment analysis of untargeted LC-MS metabolomics data from P12 maternal milk comparing control and VNMAA-treated dams. The top 25 enriched metabolite sets are shown. Dot position indicates pathway enrichment significance, plotted as −log10(p-value); dot size indicates enrichment ratio; and dot colour indicates p-value. Glycerophospholipid metabolism was among the most enriched pathways altered by VNMAA treatment. (B) Pathway enrichment analysis of untargeted LC-MS metabolomics data from P12 maternal milk comparing reciprocal FMT groups, control to VNMAA FMT and VNMAA to control FMT. The top 25 enriched metabolite sets are shown. Dot position indicates −log10(p-value), dot size indicates enrichment ratio, and dot colour indicates p-value. Glycerophospholipid metabolism was again identified among the enriched pathways, suggesting that this metabolic pathway is sensitive to maternal microbiome state and FMT-mediated remodelling. (C) Heat map of selected metabolites within the glycerophospholipid-related pathway detected in maternal milk by untargeted metabolomics. Colour intensity represents relative abundance, with darker blue indicating higher abundance. Choline is highlighted because it showed microbiome-associated variation across treatment and FMT groups. (D) Relative abundance of choline in maternal milk from control and VNMAA-treated dams, and from reciprocal FMT groups. Box plots show choline abundance in control versus VNMAA samples and in control to VNMAA FMT versus VNMAA to control FMT samples. Choline abundance was lower in VNMAA-treated dams compared with controls (p = 0.0777). In reciprocal FMT groups, choline abundance was higher in VNMAA to control FMT samples than in control to VNMAA FMT samples, although this comparison was not statistically significant (p = 0.213). These data identify maternal milk choline and glycerophospholipid metabolism as candidate microbiome-dependent pathways that may contribute to neonatal retinal vascular development.

Importantly, glycerophospholipid metabolism was also prominent in the comparison between reciprocal FMT groups, suggesting that this pathway was linked not only to antibiotic-induced dysbiosis, but also to the altered maternal microbial states generated by microbiota transfer (Figure 4B). The recurrence of this pathway across comparisons pointed to milk lipid metabolism as a potential mechanistic bridge between maternal microbiome composition and neonatal vascular development.

We therefore examined metabolites within the glycerophospholipid pathway in greater detail. Heatmap analysis revealed coordinated differences across several related metabolites, including choline, serine, methionine, total diacylglycerides, total lysophosphatidylcholine, and total phosphatidylcholine (Figure 4C). Among these, choline was of particular interest because of its known role in phospholipid metabolism, membrane biosynthesis, methyl-group metabolism, and cellular function. Although choline did not reach significance in all comparisons (Figure 4D), its position within an enriched microbiome-sensitive pathway and its biological relevance to endothelial function made it a strong candidate for functional testing.

Together, these data identify the maternal milk metabolome as a plausible mediator linking maternal microbiome composition to neonatal vascular development. Rather than implicating the neonatal gut microbiome itself as the primary driver, the metabolomic findings support a model in which the maternal microbiome shapes the biochemical composition of milk, thereby altering the metabolic signals available to the developing offspring vasculature. This led us to test whether choline, as a representative microbiome-sensitive metabolite within the glycerophospholipid pathway, could directly influence endothelial cell behaviour.

### Choline modulates endothelial viability and promotes angiogenic tubule formation

Having identified glycerophospholipid metabolism as a microbiome-sensitive pathway in maternal milk, we next asked whether a metabolite from this pathway could directly influence endothelial cell behaviour. We focused on choline because it varied across milk samples according to maternal microbial state, sits within the enriched glycerophospholipid pathway, and is central to phospholipid metabolism, membrane biosynthesis, and cellular function. We therefore tested whether choline could modulate endothelial viability and angiogenic capacity *in vitro*.

HUVECs were treated with increasing concentrations of choline, with VEGF included as a positive pro-angiogenic control and suramin as a negative control. Choline exposure altered endothelial cell viability in a concentration-dependent manner. Although VEGF produced the strongest pro-survival response, choline-treated cells showed a pattern consistent with direct modulation of endothelial cell function (Figure 5A).

**Figure 5.**
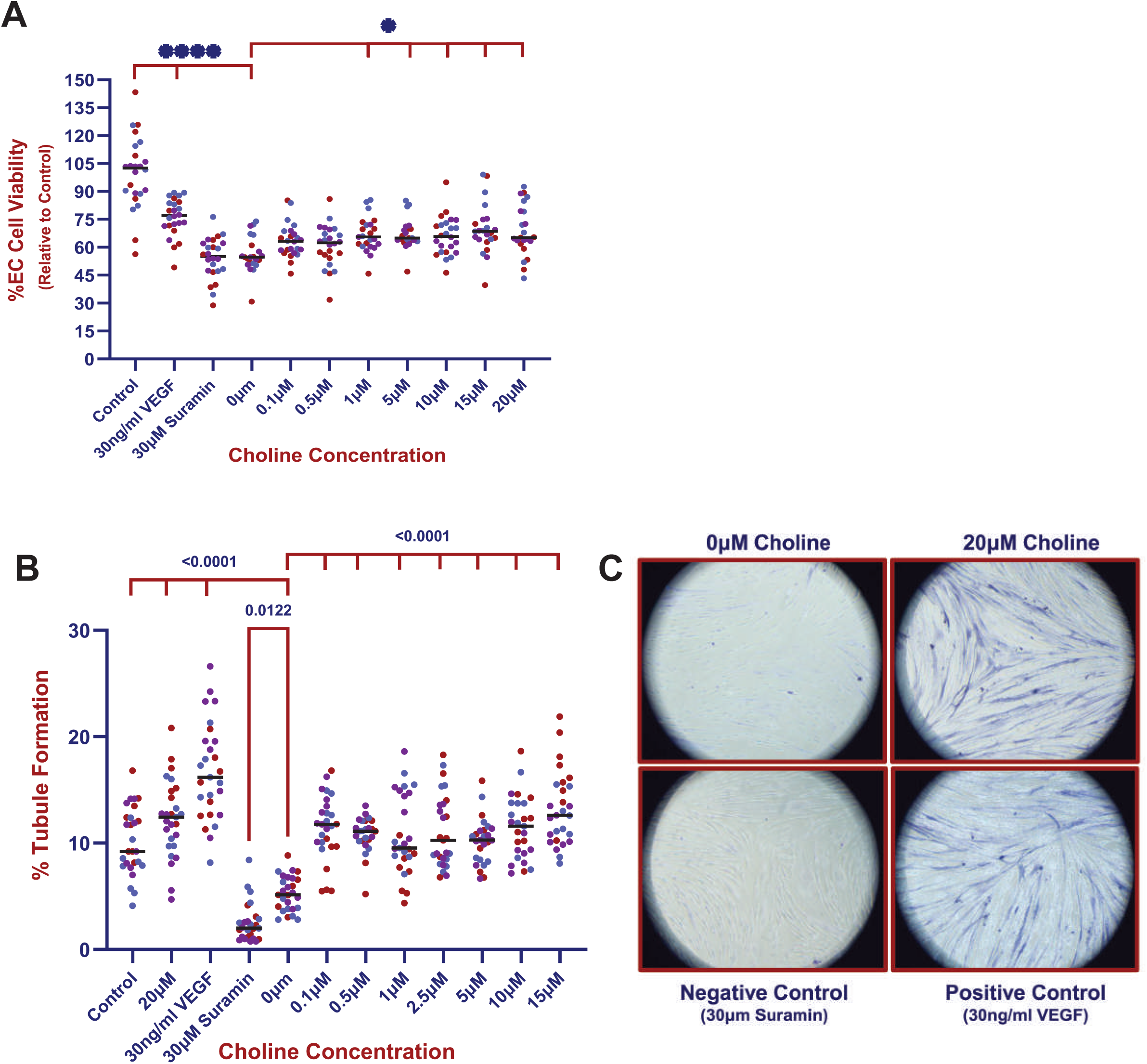
Choline promotes endothelial tubule formation without increasing endothelial cell viability. (A) Quantification of endothelial cell viability following treatment with choline. HUVECs were treated with increasing concentrations of choline, with untreated control cells, 30ng/ml VEGF, and 30µM suramin included as assay controls. Cell viability is expressed as percentage relative to control. Data are presented as superplots, where each colour represents an independent biological repeat and each point represents an individual analysed well within that biological repeat. Black horizontal lines indicate the mean. Suramin reduced cell viability compared with control. (B) Quantification of endothelial tubule formation following choline treatment. HDF/HUVEC co-cultures were treated with increasing concentrations of choline, with untreated control cells, 30ng/ml VEGF as a positive control, and 30µM suramin as a negative control. Tubule formation is expressed as percentage tubule formation. Data are presented as superplots, where each colour represents an independent biological repeat and each point represents an individual well within that biological repeat. Black horizontal lines indicate the mean. VEGF increased tubule formation compared with control, whereas suramin strongly reduced tubule formation. Choline treatment increased tubule formation compared with 0µM choline, with 20µM choline showing enhanced tubule formation relative to the untreated control condition. Indicated p-values are shown on the graph. (C) Representative images from the HDF/HUVEC tubule formation assay showing 0µM choline, 20µM choline, 30µM suramin negative control, and 30ng/ml VEGF positive control conditions. Tubule-like endothelial networks are visible in the 20µM choline and VEGF-treated conditions, whereas network formation is reduced in the 0µM choline and suramin-treated conditions. Images correspond to the tubule formation quantification shown in (B). Statistical significance is indicated on the graphs: **P < 0.01, ****P < 0.0001. Data are presented as pooled N: each colour represents an independent biological repeat, and each point represents an individual analysed well or field within that repeat. The black horizontal line indicates the group mean.

We next assessed whether choline could influence endothelial morphogenesis using a tubule formation assay. Compared with untreated controls, choline increased tubule formation, with 20µM choline producing a significant increase in network formation (Figure 5B). Representative images showed more extensive and organised tubule-like structures in choline-treated cultures compared with untreated controls, while suramin suppressed tubule formation and VEGF enhanced network formation as expected (Figure 5C).

These findings demonstrate that choline is capable of directly modulating endothelial cell behaviour *in vitro*. While choline is unlikely to be the only milk-derived metabolite contributing to the *in vivo* phenotype, these data provide functional support for the broader model that maternal microbiome-sensitive milk metabolites can influence angiogenic processes. Together with the milk metabolomics data, this establishes a mechanistic link between altered maternal microbial state, changes in milk glycerophospholipid metabolism, and endothelial behaviours required for neonatal vascular development.

### Integrated model of a microbiota–metabolite–vascular axis

Together, our findings support a model in which maternal microbial state regulates neonatal retinal angiogenesis through maternally derived metabolic signals. Maternal microbiome depletion first produced a clear vascular phenotype, delaying early retinal vascular expansion and impairing deep plexus maturation. Metagenomic profiling then showed that the most pronounced microbial remodelling occurred in dams, whereas the P12 pup caecal microbiome was comparatively limited and did not mirror the dramatic maternal shifts. This redirected the mechanistic focus toward maternal-derived factors rather than direct effects of a mature offspring gut microbiome.

Consistent with this interpretation, restoring the maternal microbiome through FMT rescued neonatal angiogenesis, while transfer of a dysbiotic microbiota shifted offspring vascular development toward an impaired phenotype. These reciprocal effects indicate that the maternal microbiome is not merely associated with neonatal vascular development but is functionally linked to it.

Milk metabolomics identified glycerophospholipid metabolism as a microbiome-sensitive pathway, with choline and related lipid metabolites emerging as candidate mediators. Functional endothelial assays further demonstrated that choline can promote angiogenic tubule formation and modulate endothelial viability, supporting a role for milk-derived metabolites in regulating angiogenic behaviour.

We therefore propose a maternal microbiota–milk metabolite–vascular axis in which maternal microbial communities shape the biochemical composition of milk, thereby altering metabolite availability to the neonate and influencing endothelial function during retinal vascular development (Figure 6). This model provides a mechanistic framework for understanding how maternal microbial ecology may contribute to early-life vascular programming.

**Figure 6.**
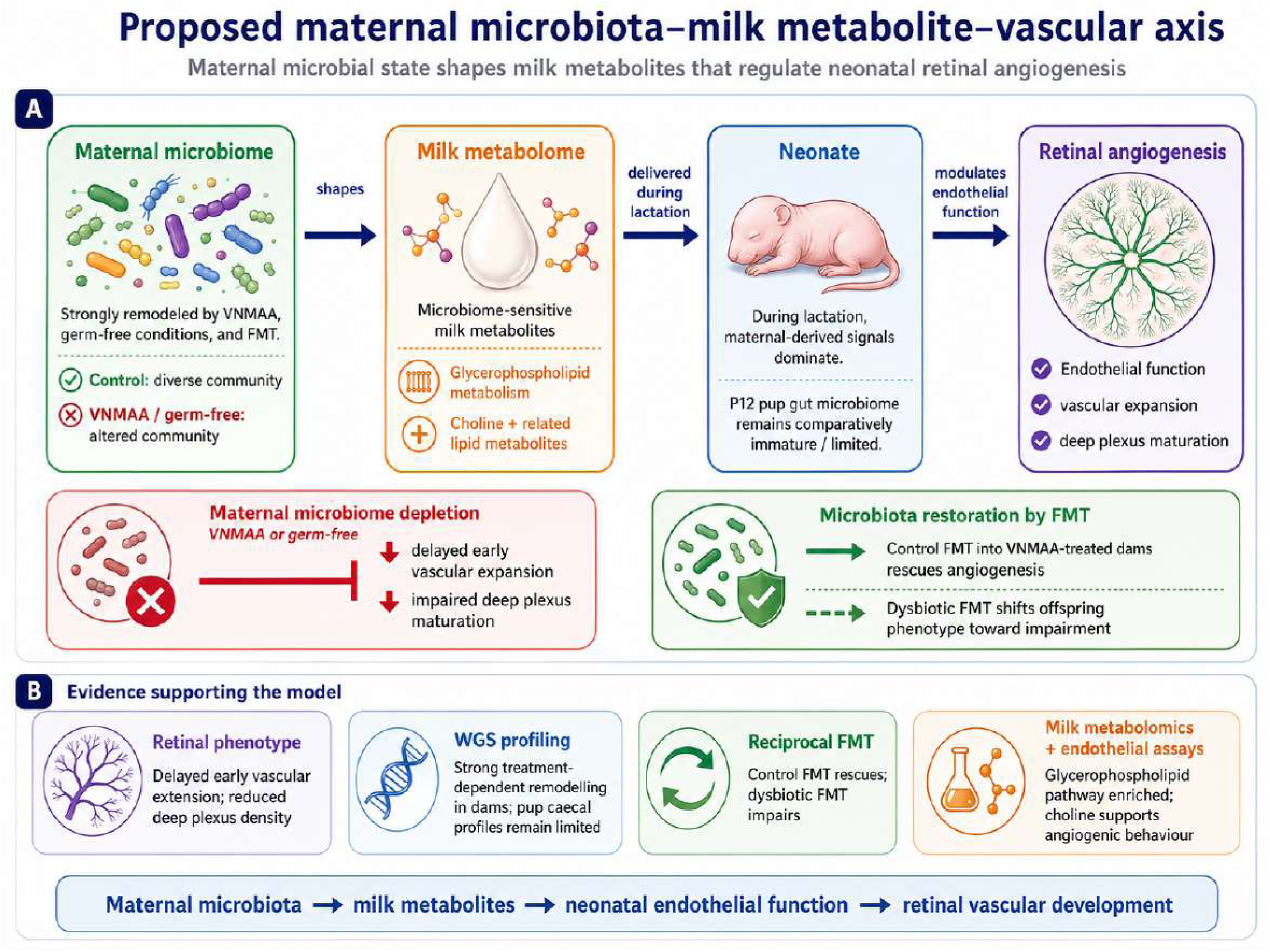
Proposed maternal microbiota–milk metabolite–vascular axis regulating neonatal retinal angiogenesis. (A) Schematic model summarising the proposed mechanism by which maternal microbial state regulates neonatal retinal vascular development. Maternal microbiome composition is altered by microbiome depletion, germ-free conditions, and reciprocal FMT. These changes are proposed to reshape the maternal milk metabolome, with glycerophospholipid metabolism and choline-related lipid metabolites emerging as microbiome-sensitive pathways. During lactation, maternally derived milk metabolites are delivered to the neonate during a developmental window in which the P12 pup gut microbiome remains comparatively immature and does not fully recapitulate the maternal microbial shifts. These milk-derived signals are proposed to modulate neonatal endothelial function, thereby influencing retinal vascular expansion and deep plexus maturation. Maternal microbiome depletion, including VNMAA treatment or germ-free status, is associated with delayed early retinal vascular expansion and impaired deep plexus maturation, whereas restoration of control microbiota by FMT into VNMAA-treated dams supports rescue of neonatal angiogenesis. Conversely, transfer of dysbiotic VNMAA-associated microbiota into control dams shifts offspring retinal vascular development toward an impaired phenotype. (B) Summary of experimental evidence supporting the model. Early-life microbiome depletion produced a retinal vascular phenotype characterised by delayed vascular extension and reduced deep plexus vascular density. Whole-genome shotgun profiling showed strong treatment-dependent remodelling of the maternal microbiome, while P12 pup caecal microbial profiles remained comparatively limited. Reciprocal FMT experiments demonstrated that the maternal microbial state is functionally linked to offspring angiogenesis, with control FMT rescuing and dysbiotic FMT impairing retinal vascular development. Milk metabolomic analysis identified glycerophospholipid metabolism as a microbiome-sensitive pathway, and endothelial assays supported a pro-angiogenic role for choline. Together, these findings support a model in which the maternal microbiota regulates milk metabolite availability, which in turn modulates neonatal endothelial function and retinal vascular development. AI image generated by ChatGPT (version 5.4).

## Discussion

This study identifies maternal microbial state as a regulator of neonatal retinal vascular development and supports a model in which this effect is mediated, at least in part, through lactational metabolic signalling. Maternal microbiome depletion delayed early radial vascular expansion at P6 and produced persistent abnormalities in vascular architecture at P12, particularly within the deep retinal plexus. These effects were recapitulated in germ-free animals and were partially reversed by restoration of the maternal microbiota through FMT, strengthening the interpretation that the phenotype reflects loss or alteration of microbial input rather than antibiotic exposure alone. By integrating retinal vascular phenotyping with maternal and pup metagenomics, milk metabolomics, and endothelial functional assays, our findings suggest that maternal microbiota can influence neonatal angiogenesis through changes in maternally derived metabolic cues.

A key feature of the phenotype was its temporal and compartment-specific nature. Microbiome depletion reduced superficial vascular extension at P6, but this difference was no longer evident by P12. At first glance, this might suggest that vascular development simply catches up over time. However, more detailed analysis of P12 vascular architecture showed that recovery of radial outgrowth did not equate to complete normalisation of retinal vascular development. Instead, microbiome-depleted animals showed persistent defects in deeper vascular compartments, including reduced deep plexus vessel density, altered descending branch density, and reduced branching complexity after normalisation to vessel density. This pattern suggests that maternal microbiome depletion does not cause a uniform arrest of angiogenesis, but rather disrupts the transition from early superficial expansion to later three-dimensional vascular plexus maturation. This distinction is biologically important because deep plexus formation requires a more complex sequence of endothelial behaviours than radial superficial outgrowth alone. After the superficial plexus expands across the retina, endothelial cells must sprout vertically, migrate into deeper retinal layers, remodel into a capillary network, and adapt to the metabolic demands of the developing neural retina ^5–8^. These processes require coordinated regulation of endothelial proliferation, migration, membrane remodelling, and branching. The sensitivity of the deep plexus in our model may therefore reflect a greater dependence on sustained metabolic support during this later phase of vascular maturation. This interpretation is consistent with the broader concept that endothelial metabolism is functionally instructive during angiogenesis, with glycolysis, lipid metabolism, and nucleotide synthesis all contributing to endothelial sprouting, proliferation, and vessel patterning^11,12,41,42^.

The metagenomic data were central to refining the mechanistic interpretation of the phenotype. Because pups are exposed to maternal microbes through birth, nursing, grooming, and shared environment, an obvious possibility was that altered pup gut colonisation directly caused the retinal vascular defects. However, the strongest treatment-associated microbial restructuring was observed in dams, whereas P12 pup caecal communities were comparatively limited and did not mirror the pronounced group-level differences seen in maternal samples. This does not exclude a contribution from the offspring microbiome, particularly at other developmental stages, mucosal sites, or through microbial metabolites not captured by caecal taxonomic profiling. Nevertheless, at the time point examined, pup caecal composition did not provide the most compelling explanation for the vascular phenotype.

This finding shifted the focus toward maternal-derived signals. The timing of mouse retinal vascular development makes this particularly relevant: retinal angiogenesis occurs largely during the early postnatal period, when pups remain dependent on maternal milk. The data therefore support a model in which maternal microbial communities influence the neonatal environment indirectly, through changes in maternal physiology and milk composition, rather than exclusively through establishment of a mature offspring gut microbiome. This interpretation aligns with previous studies showing that maternal microbiota can shape offspring immune, metabolic, and neurodevelopmental outcomes through maternally mediated mechanisms, including microbial metabolites and altered foetal or neonatal metabolite availability^17–21^.

The reciprocal FMT experiments provide important functional support for this model. Restoration of control microbiota to VNMAA-treated dams improved offspring retinal vascular density, whereas transfer of VNMAA-associated microbiota into control dams shifted the offspring phenotype toward impaired vascularisation. These reciprocal effects indicate that maternal microbial state is not only associated with neonatal vascular development but can influence the direction of the phenotype. At the same time, FMT should be interpreted as a community-level intervention. It does not identify the responsible taxa, microbial genes, or microbial products, and it may affect maternal metabolism, immune tone, and microbial metabolite production simultaneously. Its value here is therefore not in defining a single causal organism, but in demonstrating that altering the maternal microbial ecosystem is sufficient to modify offspring angiogenic outcome.

Milk metabolomics provided a plausible mechanistic bridge between maternal microbial state and neonatal endothelial function. Untargeted LC–MS analysis showed that maternal microbiome depletion and reciprocal FMT were associated with changes in milk metabolic pathways, with glycerophospholipid metabolism emerging as a prominent and recurrent signal. This pathway is biologically relevant to angiogenesis because endothelial sprouting and network formation require membrane expansion, phospholipid turnover, lipid-mediated signalling, and cellular proliferation. The broader metabolomic changes observed across amino acid, glutathione, tryptophan, fatty acid, bile acid-related, and sphingolipid-associated pathways also suggest that the maternal microbiome may reshape milk composition in a multifactorial manner. Thus, the vascular phenotype is unlikely to be driven by a single metabolite in isolation; rather, it may reflect coordinated changes in the biochemical environment available to the developing neonate.

Within this altered metabolic landscape, choline was prioritised as a functional candidate because of its position within glycerophospholipid metabolism and its established roles in phosphatidylcholine synthesis, membrane biogenesis, one-carbon metabolism, lipid transport, and cellular function^29,30^. Developmental tissues have high requirements for membrane synthesis and methyl-group metabolism, making choline availability particularly relevant during periods of rapid growth. Prior work has also linked maternal choline availability to developmental angiogenesis, with maternal choline deficiency altering endothelial proliferation, vessel number, and angiogenic gene expression in the foetal mouse hippocampus ^31^. In addition, low choline availability during pregnancy has been shown to disrupt retinal development and later visual function in mice, causing persistent defects in retinal cytoarchitecture, altered retinal progenitor cell behaviour, impaired neuronal differentiation, and abnormal adult visual performance^43^. Although this study focused primarily on retinal neurogenesis and structure rather than vascular development, it provides direct support for the relevance of maternal choline availability to visual system development. Choline insufficiency has also been shown to impair trophoblast function and *in vitro* vascularisation, altering angiogenic and inflammatory profiles and increasing apoptosis and oxidative stress in human trophoblast cultures^44^. Together, these studies strengthen the biological plausibility that maternal choline availability can influence developmental processes involving vascular growth, tissue patterning, and retinal maturation.

Our endothelial assays support choline as a biologically active candidate rather than merely a correlative metabolite. Choline promoted endothelial tubule formation and modulated endothelial viability *in vitro*, indicating that it can directly influence cellular behaviours relevant to angiogenesis. These results are consistent with the known importance of phosphatidylcholine metabolism in endothelial cells^32^ and with the wider literature linking endothelial metabolic state to angiogenic function^11,12,41,42^. However, choline should not be interpreted as the sole mediator of the *in vivo* phenotype. The milk metabolomics data point to coordinated changes across multiple lipid and metabolite classes, and the *in vitro* assays do not establish that choline is sufficient to rescue retinal angiogenesis in microbiome-depleted pups. Rather, choline provides proof-of-principle that a microbiome-sensitive milk metabolite can modulate endothelial behaviour.

Together, these findings support a maternal microbiota–milk metabolite–vascular axis. In this model, maternal microbial communities shape the biochemical composition of milk; milk-derived metabolites alter the metabolic cues available to the neonate; and these cues influence endothelial behaviours required for retinal vascular expansion and plexus maturation. The lactational component of this model is important because it extends the concept of maternal microbiome-dependent developmental programming beyond pregnancy. It suggests that maternal microbial ecology may continue to influence organ development after birth, during a period when milk is the dominant nutritional and biochemical interface between mother and offspring.

This model also raises several mechanistic questions. Maternal microbiota could alter milk glycerophospholipid and choline-related metabolites through direct microbial metabolism, modulation of host hepatic lipid handling, entero-mammary signalling, immune-mediated effects on mammary gland biology, or altered maternal nutrient utilisation. These possibilities are not mutually exclusive. Future studies combining microbial functional profiling, targeted milk metabolomics, maternal serum metabolomics, and mammary gland analysis will be needed to determine how changes in the gut microbial ecosystem are transmitted to the milk metabolome.

The findings may also have broader implications for early-life vascular programming. The retina is a useful and highly tractable model of physiological angiogenesis, but vascular development is fundamental to the maturation of many tissues. Altered endothelial growth or vascular patterning during critical windows could influence tissue oxygenation, nutrient delivery, immune trafficking, and metabolic homeostasis beyond the immediate developmental period. Within the developmental origins of health and disease framework, vascular development may therefore represent an intermediary process linking maternal exposures to later cardiometabolic or neurovascular outcomes^1–3^. However, this remains speculative, and the current study does not establish long-term functional consequences. Future work will be required to determine whether the vascular changes observed here persist, resolve, or alter susceptibility to later vascular stress.

The retinal context may also have disease relevance. Abnormal vascular development is central to paediatric retinal vascular disorders, including retinopathy of prematurity. Although the present study examines physiological retinal angiogenesis rather than oxygen-induced vascular pathology, it raises the possibility that maternal microbial state and milk metabolite composition could influence how the developing retina responds to vascular stress. Testing this directly in oxygen-induced retinopathy or other models would determine whether maternal microbiome-dependent metabolic signalling affects vascular regression, repair, or pathological neovascularisation^9,10^.

Several limitations should be acknowledged. First, while the retina is an established model of developmental angiogenesis, it may not represent all vascular beds. Mouse retinal vascularisation occurs postnatally, whereas many human vascular beds develop prenatally, and the retinal neurovascular niche has specialised cellular and metabolic features. Second, antibiotic treatment may have microbiota-independent effects on maternal metabolism, inflammation, milk production, or neonatal physiology. The inclusion of germ-free animals and reciprocal FMT mitigates this concern but does not eliminate it. Defined gnotobiotic colonisation or selective microbial add-back experiments would provide stronger mechanistic resolution.

Third, the pup caecal microbiome data should be interpreted cautiously. The comparatively limited P12 pup profiles suggest that mature pup caecal colonisation is unlikely to be the dominant explanatory factor at this time point, but they do not rule out contributions from earlier colonisation events, mucosal-associated communities, microbial products, or circulating microbial metabolites. Fourth, milk metabolomics was performed at a single lactational time point. Because milk composition changes dynamically across lactation, the metabolites most relevant to P6 vascular extension may differ from those influencing P12 deep plexus maturation. Longitudinal milk profiling would help define the critical windows of metabolic exposure.

Fifth, the metabolomics data are discovery-based. Targeted quantification of choline, phosphatidylcholine species, lysophosphatidylcholines, diacylglycerides, and related metabolites will be needed to validate the pathway-level findings and establish dose relationships. Sixth, choline was tested *in vitro* but not yet *in vivo*. Supplementation or rescue experiments in microbiome-depleted dams or pups would be required to determine whether choline is sufficient to restore retinal angiogenesis, and whether its effects depend on timing, dose, or interaction with other milk metabolites. Finally, the HUVEC and HDF/HUVEC co-culture systems provide useful functional assays but do not fully recapitulate neonatal mouse retinal endothelial biology. Future studies using primary retinal endothelial cells, retinal explants, endothelial-specific metabolic readouts, or *in vivo* supplementation approaches would strengthen mechanistic relevance.

In summary, this study identifies maternal microbial state as a regulator of neonatal retinal angiogenesis and supports lactational metabolic signalling as a plausible mechanism. Maternal microbiome depletion delayed early retinal vascular expansion and produced persistent defects in deep plexus maturation, descending branch formation, and vascular branching complexity, while restoration of the maternal microbiota improved the vascular phenotype. Metagenomic profiling indicated that the maternal microbiome was the dominant remodelled microbial compartment, and milk metabolomics implicated glycerophospholipid metabolism as a microbiome-sensitive pathway. Choline emerged as a functional candidate metabolite capable of modulating endothelial behaviour *in vitro*. Together, these findings expand the developmental influence of the maternal microbiome to include regulation of neonatal vascular development and provide a framework for future studies defining the microbial, metabolic, and endothelial mechanisms linking maternal microbial ecology to offspring vascular health.

## Acknowledgments

SAD and SDR gratefully acknowledge the support of the Biotechnology and Biological Sciences Research Council (BBSRC); this research was funded by the BBSRC Institute Strategic Programme Food Microbiome and Health BB/X011054/1 and its constituent project(s) BBS/E/QU/230001A, BBS/E/QU/230001B, BBS/E/QU/230001C.

We would like to thank the Quadram Institute Bioimaging Facility and the Quadram Institute Sequencing Facility for their support with imaging and sequencing respectively. We would also like to thank the Disease Modelling Unit Facility at the University of East Anglia for their assistance with the animals and a special acknowledgment to Arlaine Brion at QIB Germ-free Facility for her help with providing the germ-free animals in this study. We are grateful to Professor Lindsay Hall for her support of this study and for enabling Raymond Kiu’s contribution to the metagenomic sequencing and taxonomic profiling analyses.

## Author contributions

Conceptualisation: SAD; Formal analyses: SAD; Bioinformatics analyses: RK; Investigation: SAD; Resources: SDR; Review and editing: SAD, SDR; Visualisation: SAD, SDR; Supervision: SDR; Funding acquisition: SDR.

## References

1. Barker, D.J., and Osmond, C. (1986). Infant mortality, childhood nutrition, and ischaemic heart disease in England and Wales. Lancet 1, 1077–1081. 10.1016/s0140-6736(86)91340-1.

2. Barker, D.J. (1990). The fetal and infant origins of adult disease. BMJ 301, 1111. 10.1136/bmj.301.6761.1111.

3. Gluckman, P.D., and Hanson, M.A. (2004). Living with the past: evolution, development, and patterns of disease. Science 305, 1733–1736. 10.1126/science.1095292.

4. Potente, M., Gerhardt, H., and Carmeliet, P. (2011). Basic and therapeutic aspects of angiogenesis. Cell 146, 873–887. 10.1016/j.cell.2011.08.039.

5. Fruttiger, M. (2002). Development of the mouse retinal vasculature: angiogenesis versus vasculogenesis. Invest Ophthalmol Vis Sci 43, 522–527.

6. Gariano, R.F. (2003). Cellular mechanisms in retinal vascular development. Prog Retin Eye Res 22, 295–306. 10.1016/s1350-9462(02)00062-9.

7. Stahl, A., Connor, K.M., Sapieha, P., Chen, J., Dennison, R.J., Krah, N.M., Seaward, M.R., Willett, K.L., Aderman, C.M., Guerin, K.I., et al. (2010). The mouse retina as an angiogenesis model. Invest Ophthalmol Vis Sci 51, 2813–2826. 10.1167/iovs.10-5176.

8. Fruttiger, M. (2007). Development of the retinal vasculature. Angiogenesis 10, 77–88. 10.1007/s10456-007-9065-1.

9. Smith, L.E., Wesolowski, E., McLellan, A., Kostyk, S.K., D’Amato, R., Sullivan, R., and D’Amore, P.A. (1994). Oxygen-induced retinopathy in the mouse. Invest Ophthalmol Vis Sci 35, 101–111.

10. Connor, K.M., Krah, N.M., Dennison, R.J., Aderman, C.M., Chen, J., Guerin, K.I., Sapieha, P., Stahl, A., Willett, K.L., and Smith, L.E. (2009). Quantification of oxygen-induced retinopathy in the mouse: a model of vessel loss, vessel regrowth and pathological angiogenesis. Nat Protoc 4, 1565–1573. 10.1038/nprot.2009.187.

11. De Bock, K., Georgiadou, M., Schoors, S., Kuchnio, A., Wong, B.W., Cantelmo, A.R., Quaegebeur, A., Ghesquiere, B., Cauwenberghs, S., Eelen, G., et al. (2013). Role of PFKFB3-driven glycolysis in vessel sprouting. Cell 154, 651–663. 10.1016/j.cell.2013.06.037.

12. Eelen, G., de Zeeuw, P., Treps, L., Harjes, U., Wong, B.W., and Carmeliet, P. (2018). Endothelial Cell Metabolism. Physiol Rev 98, 3–58. 10.1152/physrev.00001.2017.

13. Yatsunenko, T., Rey, F.E., Manary, M.J., Trehan, I., Dominguez-Bello, M.G., Contreras, M., Magris, M., Hidalgo, G., Baldassano, R.N., Anokhin, A.P., et al. (2012). Human gut microbiome viewed across age and geography. Nature 486, 222–227. 10.1038/nature11053.

14. Backhed, F., Roswall, J., Peng, Y., Feng, Q., Jia, H., Kovatcheva-Datchary, P., Li, Y., Xia, Y., Xie, H., Zhong, H., et al. (2015). Dynamics and Stabilization of the Human Gut Microbiome during the First Year of Life. Cell Host Microbe 17, 690–703. 10.1016/j.chom.2015.04.004.

15. Bokulich, N.A., Chung, J., Battaglia, T., Henderson, N., Jay, M., Li, H., A, D.L., Wu, F., Perez-Perez, G.I., Chen, Y., et al. (2016). Antibiotics, birth mode, and diet shape microbiome maturation during early life. Sci Transl Med 8, 343ra382. 10.1126/scitranslmed.aad7121.

16. Bogaert, D., van Beveren, G.J., de Koff, E.M., Lusarreta Parga, P., Balcazar Lopez, C.E., Koppensteiner, L., Clerc, M., Hasrat, R., Arp, K., Chu, M., et al. (2023). Mother-to-infant microbiota transmission and infant microbiota development across multiple body sites. Cell Host Microbe 31, 447–460 e446. 10.1016/j.chom.2023.01.018.

17. Gomez de Aguero, M., Ganal-Vonarburg, S.C., Fuhrer, T., Rupp, S., Uchimura, Y., Li, H., Steinert, A., Heikenwalder, M., Hapfelmeier, S., Sauer, U., et al. (2016). The maternal microbiota drives early postnatal innate immune development. Science 351, 1296–1302. 10.1126/science.aad2571.

18. Kimura, I., Miyamoto, J., Ohue-Kitano, R., Watanabe, K., Yamada, T., Onuki, M., Aoki, R., Isobe, Y., Kashihara, D., Inoue, D., et al. (2020). Maternal gut microbiota in pregnancy influences offspring metabolic phenotype in mice. Science 367. 10.1126/science.aaw8429.

19. Vuong, H.E., Pronovost, G.N., Williams, D.W., Coley, E.J.L., Siegler, E.L., Qiu, A., Kazantsev, M., Wilson, C.J., Rendon, T., and Hsiao, E.Y. (2020). The maternal microbiome modulates fetal neurodevelopment in mice. Nature 586, 281–286. 10.1038/s41586-020-2745-3.

20. Pessa-Morikawa, T., Husso, A., Karkkainen, O., Koistinen, V., Hanhineva, K., Iivanainen, A., and Niku, M. (2022). Maternal microbiota-derived metabolic profile in fetal murine intestine, brain and placenta. BMC Microbiol 22, 46. 10.1186/s12866-022-02457-6.

21. Husso, A., Pessa-Morikawa, T., Koistinen, V.M., Karkkainen, O., Kwon, H.N., Lahti, L., Iivanainen, A., Hanhineva, K., and Niku, M. (2023). Impacts of maternal microbiota and microbial metabolites on fetal intestine, brain, and placenta. BMC Biol 21, 207. 10.1186/s12915-023-01709-9.

22. Ballard, O., and Morrow, A.L. (2013). Human milk composition: nutrients and bioactive factors. Pediatr Clin North Am 60, 49–74. 10.1016/j.pcl.2012.10.002.

23. Sundekilde, U.K., Downey, E., O’Mahony, J.A., O’Shea, C.A., Ryan, C.A., Kelly, A.L., and Bertram, H.C. (2016). The Effect of Gestational and Lactational Age on the Human Milk Metabolome. Nutrients 8. 10.3390/nu8050304.

24. Alexandre-Gouabau, M.C., Moyon, T., David-Sochard, A., Fenaille, F., Cholet, S., Royer, A.L., Guitton, Y., Billard, H., Darmaun, D., Roze, J.C., and Boquien, C.Y. (2019). Comprehensive Preterm Breast Milk Metabotype Associated with Optimal Infant Early Growth Pattern. Nutrients 11. 10.3390/nu11030528.

25. Li, T., Samuel, T.M., Zhu, Z., Howell, B., Cho, S., Baluyot, K., Hazlett, H., Elison, J.T., Wu, D., Hauser, J., et al. (2022). Joint analyses of human milk fatty acids, phospholipids, and choline in association with cognition and temperament traits during the first 6 months of life. Front Nutr 9, 919769. 10.3389/fnut.2022.919769.

26. Ingvordsen Lindahl, I.E., Artegoitia, V.M., Downey, E., O’Mahony, J.A., O’Shea, C.A., Ryan, C.A., Kelly, A.L., Bertram, H.C., and Sundekilde, U.K. (2019). Quantification of Human Milk Phospholipids: the Effect of Gestational and Lactational Age on Phospholipid Composition. Nutrients 11. 10.3390/nu11020222.

27. Isganaitis, E., Venditti, S., Matthews, T.J., Lerin, C., Demerath, E.W., and Fields, D.A. (2019). Maternal obesity and the human milk metabolome: associations with infant body composition and postnatal weight gain. Am J Clin Nutr 110, 111–120. 10.1093/ajcn/nqy334.

28. Saben, J.L., Sims, C.R., Piccolo, B.D., and Andres, A. (2020). Maternal adiposity alters the human milk metabolome: associations between nonglucose monosaccharides and infant adiposity. Am J Clin Nutr 112, 1228–1239. 10.1093/ajcn/nqaa216.

29. Zeisel, S.H. (2006). Choline: critical role during fetal development and dietary requirements in adults. Annu Rev Nutr 26, 229–250. 10.1146/annurev.nutr.26.061505.111156.

30. Sanders, L.M., and Zeisel, S.H. (2007). Choline: Dietary Requirements and Role in Brain Development. Nutr Today 42, 181–186. 10.1097/01.NT.0000286155.55343.fa.

31. Mehedint, M.G., Craciunescu, C.N., and Zeisel, S.H. (2010). Maternal dietary choline deficiency alters angiogenesis in fetal mouse hippocampus. Proc Natl Acad Sci U S A 107, 12834–12839. 10.1073/pnas.0914328107.

32. Wong, J.T., Chan, M., Lee, D., Jiang, J.Y., Skrzypczak, M., and Choy, P.C. (2000). Phosphatidylcholine metabolism in human endothelial cells: modulation by phosphocholine. Mol Cell Biochem 207, 95–100. 10.1023/a:1007054601256.

33. Costello, S.P., Conlon, M.A., Vuaran, M.S., Roberts-Thomson, I.C., and Andrews, J.M. (2015). Faecal microbiota transplant for recurrent Clostridium difficile infection using long-term frozen stool is effective: clinical efficacy and bacterial viability data. Aliment Pharmacol Ther 42, 1011–1018. 10.1111/apt.13366.

34. Benwell, C.J., Taylor, J., and Robinson, S.D. (2021). Endothelial neuropilin-2 influences angiogenesis by regulating actin pattern development and alpha5-integrin-p-FAK complex recruitment to assembling adhesion sites. FASEB J 35, e21679. 10.1096/fj.202100286R.

35. McKee, A.M., Kirkup, B.M., Madgwick, M., Fowler, W.J., Price, C.A., Dreger, S.A., Ansorge, R., Makin, K.A., Caim, S., Le Gall, G., et al. (2021). Antibiotic-induced disturbances of the gut microbiota result in accelerated breast tumor growth. iScience 24, 103012. 10.1016/j.isci.2021.103012.

36. Chen, S., Zhou, Y., Chen, Y., and Gu, J. (2018). fastp: an ultra-fast all-in-one FASTQ preprocessor. Bioinformatics 34, i884–i890. 10.1093/bioinformatics/bty560.

37. Wood, D.E., Lu, J., and Langmead, B. (2019). Improved metagenomic analysis with Kraken 2. Genome Biol 20, 257. 10.1186/s13059-019-1891-0.

38. Lu, J., Breitwieser, F.P., Thielen, P., and Salzberg, S.L. (2017). Bracken: estimating species abundance in metagenomics data. PeerJ Comput Sci 3. 10.7717/peerj-cs.104.

39. Blanco-Miguez, A., Beghini, F., Cumbo, F., McIver, L.J., Thompson, K.N., Zolfo, M., Manghi, P., Dubois, L., Huang, K.D., Thomas, A.M., et al. (2023). Extending and improving metagenomic taxonomic profiling with uncharacterized species using MetaPhlAn 4. Nat Biotechnol 41, 1633–1644. 10.1038/s41587-023-01688-w.

40. Beghini, F., McIver, L.J., Blanco-Miguez, A., Dubois, L., Asnicar, F., Maharjan, S., Mailyan, A., Manghi, P., Scholz, M., Thomas, A.M., et al. (2021). Integrating taxonomic, functional, and strain-level profiling of diverse microbial communities with bioBakery 3. Elife 10. 10.7554/eLife.65088.

41. Schoors, S., Bruning, U., Missiaen, R., Queiroz, K.C., Borgers, G., Elia, I., Zecchin, A., Cantelmo, A.R., Christen, S., Goveia, J., et al. (2015). Fatty acid carbon is essential for dNTP synthesis in endothelial cells. Nature 520, 192–197. 10.1038/nature14362.

42. Li, X., Sun, X., and Carmeliet, P. (2019). Hallmarks of Endothelial Cell Metabolism in Health and Disease. Cell Metab 30, 414–433. 10.1016/j.cmet.2019.08.011.

43. Trujillo-Gonzalez, I., Friday, W.B., Munson, C.A., Bachleda, A., Weiss, E.R., Alam, N.M., Sha, W., Zeisel, S.H., and Surzenko, N. (2019). Low availability of choline in utero disrupts development and function of the retina. FASEB J 33, 9194–9209. 10.1096/fj.201900444R.

44. Jiang, X., Jones, S., Andrew, B.Y., Ganti, A., Malysheva, O.V., Giallourou, N., Brannon, P.M., Roberson, M.S., and Caudill, M.A. (2014). Choline inadequacy impairs trophoblast function and vascularization in cultured human placental trophoblasts. J Cell Physiol 229, 1016–1027. 10.1002/jcp.24526.

